# Inflammatory cellular response to mechanical ventilation in elastase-induced experimental emphysema: role of pre-existing alveolar macrophages infiltration

**DOI:** 10.1101/205989

**Authors:** Anahita Rouze, Guillaume Voiriot, Elise Guivarch, Françoise Roux, Jeanne Tran Van Nhieu, Daniel Isabey, Laurent Brochard, Bernard Maitre, Armand Mekontso-Dessap, Jorge Boczkowski

**Author notes:** CORRESPONDING AUTHOR: Dr Anahita Rouzé Centre de Réanimation, CHU Lille, Boulevard du Pr Leclercq, F-59000 Lille, France. Email adresses of all co-authors.

## Abstract

**Background:** An excessive pulmonary inflammatory response could explain the poor prognosis of chronic obstructive pulmonary disease (COPD) patients submitted to invasive mechanical ventilation. The aim of this study was to evaluate the response to normal tidal volume (Vt) mechanical ventilation in a murine model of pulmonary emphysema, which represents the alveolar component of COPD. In this model, two time points associated with different levels of lung inflammation but similar lung destruction, were analyzed.

**Methods:** C57BL/6 mice received a tracheal instillation of 5 IU of porcine pancreatic elastase (Elastase mice) or the same volume of saline (Saline mice). Fourteen (D14) and 21 (D21) days after instillation, mice were anesthetized, intubated, and either mechanically ventilated (MV) with a normal Vt (8 mL/kg) or maintained on spontaneous ventilation (SV) during two hours. We analyzed respiratory mechanics, emphysema degree (mean chord length by lung histological analysis), and lung inflammation (bronchoalveolar lavage (BAL) cellularity, proportion and activation of total lung inflammatory cells by flow cytometry).

**Results:** As compared with Saline mice, Elastase mice showed a similarly increased mean chord length and pulmonary compliance at D14 and D21, while BAL cellularity was comparable between groups. Lung mechanics was similarly altered during mechanical ventilation in Elastase and Saline mice. Activated alveolar macrophages CD11b*mid* were present in lung parenchyma in both Elastase SV mice and Elastase MV mice at D14 but were absent at D21 and in Saline mice, indicating an inflammatory state with elastase at D14 only. At D14, Elastase MV mice showed a significant increase in percentage of neutrophils concomitant with a decrease in percentage of alveolar macrophages in total lung, as compared with Elastase SV mice. Furthermore, alveolar macrophages of Elastase MV mice at D14 overexpressed Gr1, and monocytes showed a trend to overexpression of CD62L, compared with Elastase SV mice.

**Conclusions:** In an elastase-induced model of pulmonary emphysema, normal Vt mechanical ventilation produced an increase in the proportion of pulmonary neutrophils, and an activation of alveolar macrophages and pulmonary monocytes. This response was observed only when the emphysema model showed an underlying inflammation (D14), reflected by the presence of activated alveolar macrophages CD11b*mid*.

## MAIN TEXT

### BACKGROUND

Chronic obstructive pulmonary disease (COPD) is characterized by persistent airflow limitation, associated with an excessive inflammatory response to noxious particles or gases in the airways and the lung (1). Invasive mechanical ventilation may lead to a higher mortality rate in this population (2), and has been recognized as an independent risk factor for mortality among COPD patients admitted to intensive care units (ICU) with acute respiratory failure (3).

Numerous experimental and clinical studies have reported the concept of ventilator-induced lung injury (VILI) (4–8). The use of high tidal volumes (Vt) is one of its main contributors, especially leading to an acute inflammation secondary to lung overdistension, and known as “biotrauma” (9–13). Normal Vt, close to 8 mL/kg may also lead to pulmonary inflammation (12,14,15). Furthermore, a preexisting lung inflammatory process could aggravate the inflammatory response to mechanical ventilation (12,16).

Since chronic airway and lung inflammation plays a major role in the pathogenesis of COPD, and its alveolar component, emphysema (17–19), we hypothesized that mechanical ventilation may aggravate preexisting pulmonary inflammation in emphysematous lungs. This phenomenon could explain, at least in part, the detrimental effect of invasive mechanical ventilation in COPD patients. The aim of our study was therefore to evaluate the inflammatory response during two hours of normal Vt mechanical ventilation in a murine model of pulmonary emphysema induced by elastase (20). This model is characterized by an early and transient alveolar infiltration by macrophages (21–23). Thereby, in order to examine the effects of pre-existing alveolar macrophages infiltration in the inflammatory response to mechanical ventilation, animals were examined at days 14 and 21 after elastase instillation, two time points associated with similar lung destruction, but the presence and the absence of macrophages infiltration, respectively.

## METHODS

Additional details on the methods are provided in an additional file.

### Animal model

All the experiments were performed in accordance with the official regulations of the French Ministry of Agriculture and the US National Institute of Health guidelines for the experimental use of animals; and were approved by the local institutional animal care and use committee. Six-weeks-old male C57BL/6 mice (Janvier Labs, Le Genest Saint-Isle, France) received the instillation of either 5 IU of porcine pancreatic elastase (Elastin Products, Owensville, MO, USA) (Elastase mice), or 50 μl of saline (Saline mice) into the trachea (23). Mice were then subjected to subsequent ventilation at two time points, 14 and 21 days after instillation.

### Mechanical ventilation

Mice were anesthetized and intubated. Mice in mechanical ventilation (MV) group were ventilated for two hours by means of a computer-driven small-animal ventilator (*FlexiVent*, Scireq, Montreal, Canada) as follows: Vt = 8 μL/g of body weight, respiratory rate = 180 /min, end-expiratory pressure = 1.5 cmH_2_O, and fraction of inspired oxygen = 0.4 - 0.6 (24). A control group (SV) consisted of anesthetized, intubated mice, maintained on spontaneous ventilation for two hours.

### Experimental design

The experimental design included four groups, at two distinct time points from tracheal instillation (D14 and D21) (figure 1): Saline SV (saline instillation followed by spontaneous ventilation), Elastase SV (elastase instillation followed by spontaneous ventilation), Saline MV (saline instillation followed by mechanical ventilation), and Elastase MV (elastase instillation followed by mechanical ventilation).

**Figure 1.**
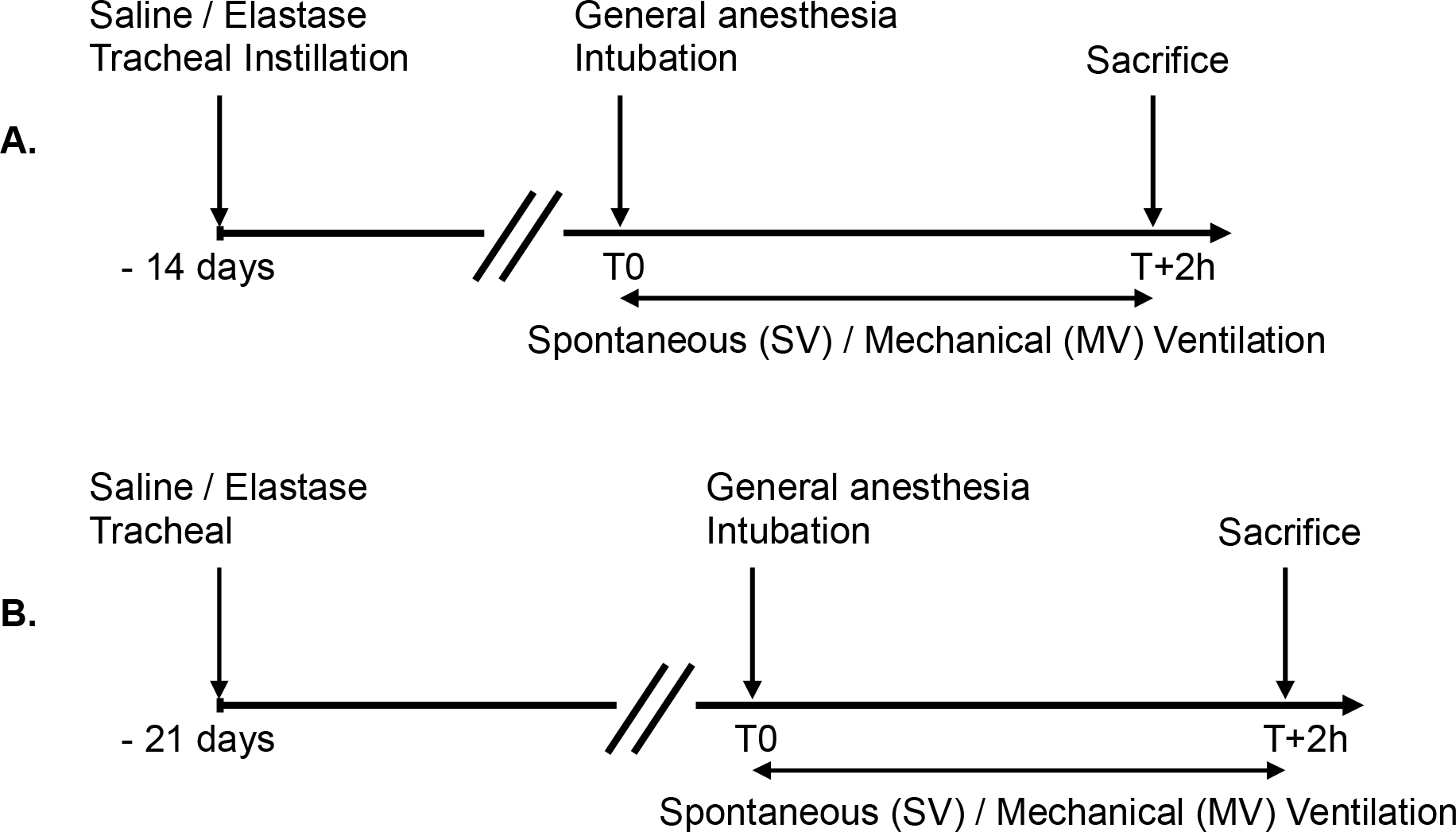
Experimental design. C57BL/6 mice received a tracheal instillation of either elastase (Elastase mice) or saline (Saline mice). After 14 (A) or 21 (B) days, mice were anesthetized, intubated and subjected to either spontaneous ventilation (SV mice) or mechanical ventilation (MV mice), then sacrificed after two hours.

### Respiratory mechanics

The *flexiVent* ventilator was used for continuous measurement of mean and peak airway pressures (Ppeak, Pmean) and to explore the respiratory mechanics using different waveforms (24). Single frequency forced oscillation techniques were assessed at initiation of mechanical ventilation, before (H0) and after (H0’) volume history standardization, and then repeated hourly (H1 and H2), for calculation of respiratory system dynamic resistance and compliance.

### Specimen collection

After sacrifice, bronchoalveolar lavage (BAL) was performed, lungs were collected for either fixation (4% paraformaldehide) and paraffin embedding, or flow cytometric analysis.

### Morphometric analysis

Sections of 5 μm thickness of the left lung were stained with hematoxylin and eosin. Fifteen digital photomicrographs were acquired at x200 magnification in a systematic fashion (Axioplan 2 microscope equipped with an MRc digital color camera (Zeiss, Oberkochen, Germany)). Emphysema was quantified by measurement of alveolar diameters with an image analysis software (ImageJ, NIH, Bethesda, USA). This automated analysis was made vertically and horizontally on each photomicrograph. The mean chord length of alveoli was obtained by averaging those measurements (23).

### Bronchoalveolar lavage

The total cell count of BAL was determined using a Malassez hemocytometer (Hycor biomedical, Indianapolis, IN, USA). Differential cell counts were done on cytocentrifuge preparations (Cytospin3; Shandon Scientific, Cheshire, UK) stained with Diff-Quick stain (Baxter Diagnostics, McGaw Park, IL, USA).

### Flow cytometric analysis

Mechanical disruption followed by enzymatic digestion of murine lungs were performed (11,25). Total lung cell suspensions were obtained, stained with fluorochrome-conjugated anti-mouse antibodies for CD11b, CD11c, Gr1 (Ly6C/G), F4/80 and CD62L (L-selectin) or appropriate isotype-matched controls, and analysed using a 7 channel cytometer (CyAn ADP Analyser, Beckman Coulter, Brea, USA) with Summit software (Summit v4.3, Dako, Cambridge, UK). Three inflammatory cell populations in murine lung were identified (figure 2). CD11b−, CD11c+ phenotype, with high autofluorescence defined alveolar macrophages (figure 2C) (25). Monocytes and neutrophils were identified as CD11b+, CD11c− cells but differed especially in their granularity (side-scatter, SS), and F4/80 and Gr1 expression (11,12). High SS and F4/80−, Gr1+ phenotype defined neutrophils, whereas low SS, and F4/80+, Gr1*mid* phenotype characterized monocytes (figures 2D, 2E, 2F). Cells activation state was assessed using expression of CD62L and CD11b adhesion molecule, as well as Gr1. All flow cytometric results were presented as relative values, called percentage of gated cells (26).

**Figure 2.**
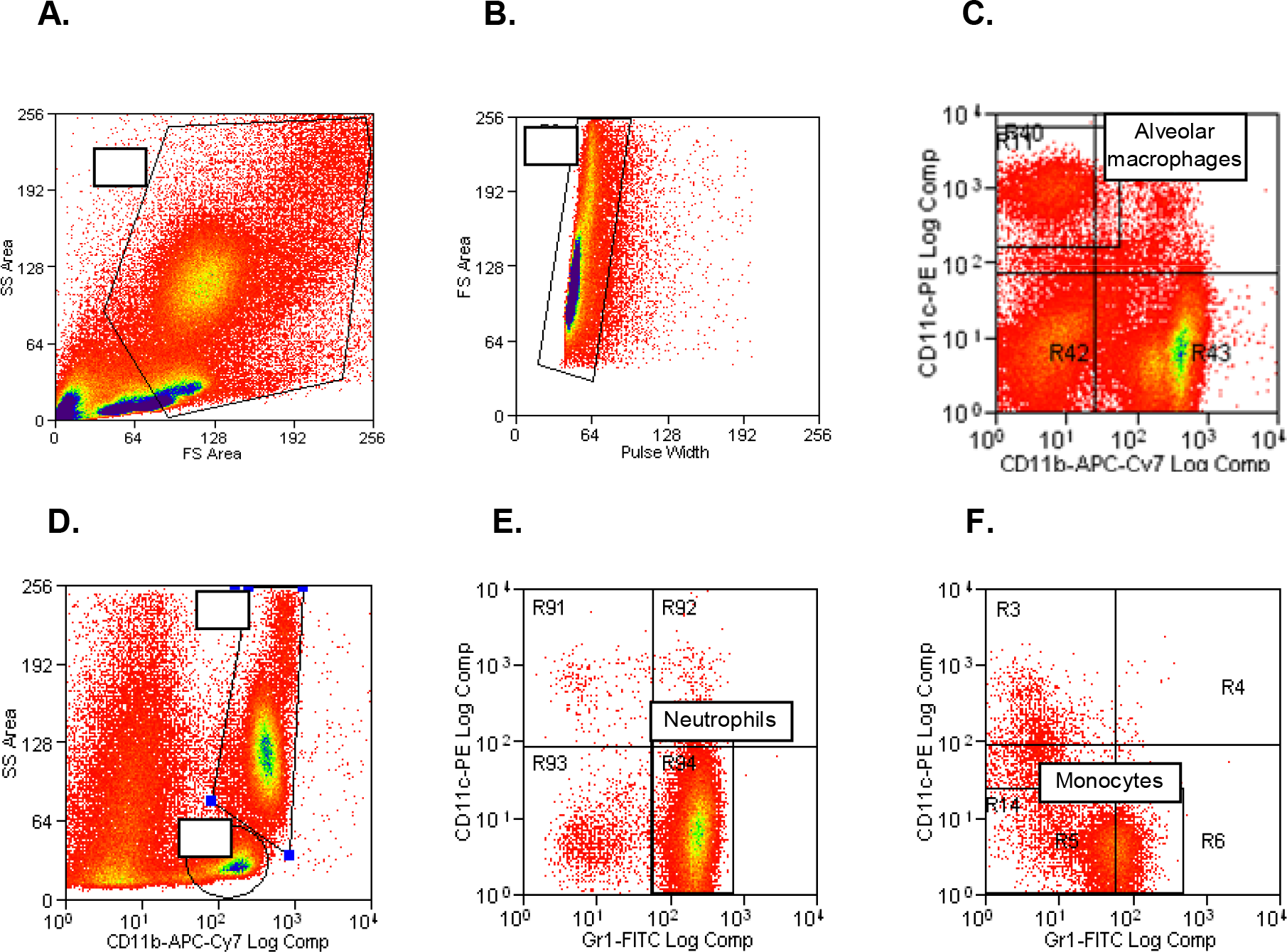
Flow cytometric analysis of different inflammatory cell populations in mouse lungs. **A.** Dot-plot showing side-scatter (SS) versus forward-scatter (FS) within total cell population. Cellular debris and lymphocyte were excluded from the analysis. Myeloid cells were gated (G1) on compatible size (FS) and granularity (SS). **B.** Dot-plot showing FS versus pulse width within G1 gated events. Doublets and triplets were subtracted from cell analysis through a second gate (G2) defined on pulse width, which distinguished cell aggregates from unique events (called singlets). From there, only combined G1 and G2 gated events were analysed. **C.** Dot-plot showing CD11c versus CD11b expression within combined G1 and G2 gated events. Alveolar macrophages were identified by CD11c+ CD11b− phenotype. Autofluorescence level was high in this cell population. **D.** Dot-plot SS versus CD11b expression within combined G1 and G2 gated events. G3 gated CD11b+ cells with neutrophils compatible granularity, G4 gated CD11b+ cells with monocytes compatible granularity. **E.** Dot plot CD11c versus Gr1 expression within combined G1, G2 and G3 gated events. Neutrophils were defined as CD11c− Gr1+ cells. **F.** Dot plot CD11c versus Gr1 expression within combined G1, G2 and G4 gated events. Monocytes were defined as CD11c− Gr1*mid* cells.

### Statistical analysis

Data were analyzed using SPSS Base 16.0 statistical software (SPSS Inc, Chicago, IL, USA). Continuous data were expressed as median ± interquartile range. Independent sample were compared using Kruskal-Wallis test followed by pairwise Mann-Whitney test, with correction for multiple testing by Benjamini-Hochberg false discovery rate. Two-tailed p values smaller than 0.10 and 0.05 were considered marginally significant and significant respectively.

## RESULTS

Additional tables and figures are provided in additional file.

### Elastase-induced emphysema model

Morphometric analysis showed a marked increase in mean chord length in Elastase mice (at both D14 and D21) as compared to Saline mice (Figure 3, Table S3 in additional file). Elastase mice (at both D14 and D21) also exhibited higher values of respiratory system dynamic compliance at the beginning of mechanical ventilation (after volume history standardisation), as compared to Saline mice (Table 1, Figure 4).

**Figure 3.**
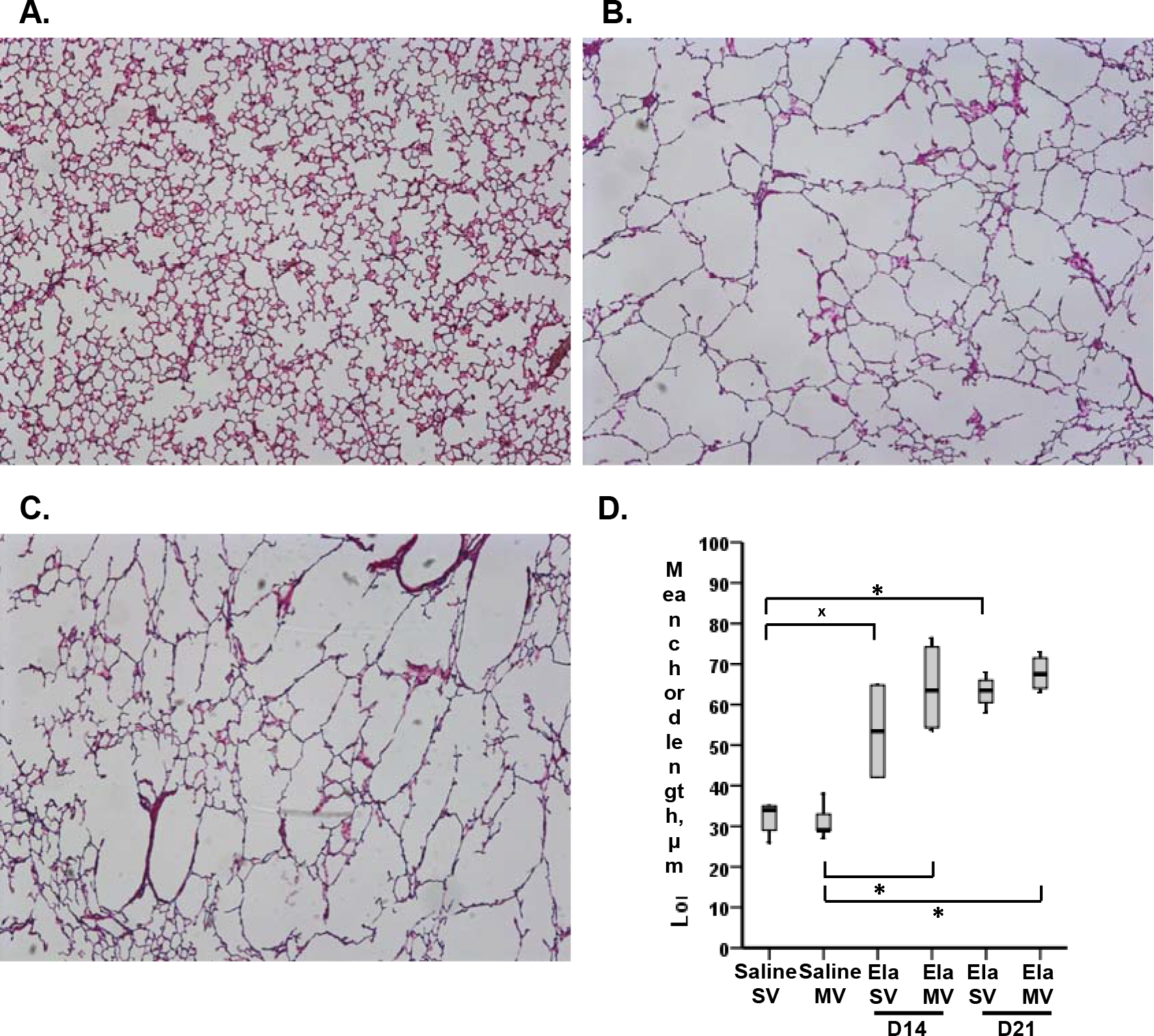
Morphometric analysis. **A, B, C.** Representative photomicrographs (50-fold magnification) of histological slides of murine lungs. Hematoxylin-eosin staining. Saline SV mice, 21 days after tracheal instillation of saline (A), Elastase SV mice, D14 (B), Elastase SV mice, D21 (C). **D.** Mean chord length of alveoli in SV and MV mice, 21 days after saline tracheal instillation, 14 and 21 days after elastase tracheal instillation. Values are expressed as medians ± interquartile range. n = 4-5 animals /group. ^±^ and * denote Benjamini-Hochberg corrected p value <0.10 (marginally significant) and <0.05 respectively for the following Mann-Whitney pairwise comparisons (following Kruskal-Wallis test): Saline SV *vs* Saline MV, Saline SV *vs* Ela SV D14, Saline SV *vs* Ela SV D21, Saline MV *vs* Ela MV D14, Saline MV *vs* Ela MV D21, Ela SV *vs* Ela MV D14, Ela SV *vs* Ela MV D21, Ela SV D14 *vs* Ela SV D21, Ela MV D14 *vs* Ela MV D21. *Definition of abbreviations*: SV, spontaneous ventilation; MV, mechanical ventilation; Ela, elastase tracheal instillation.

**TABLE 1.**
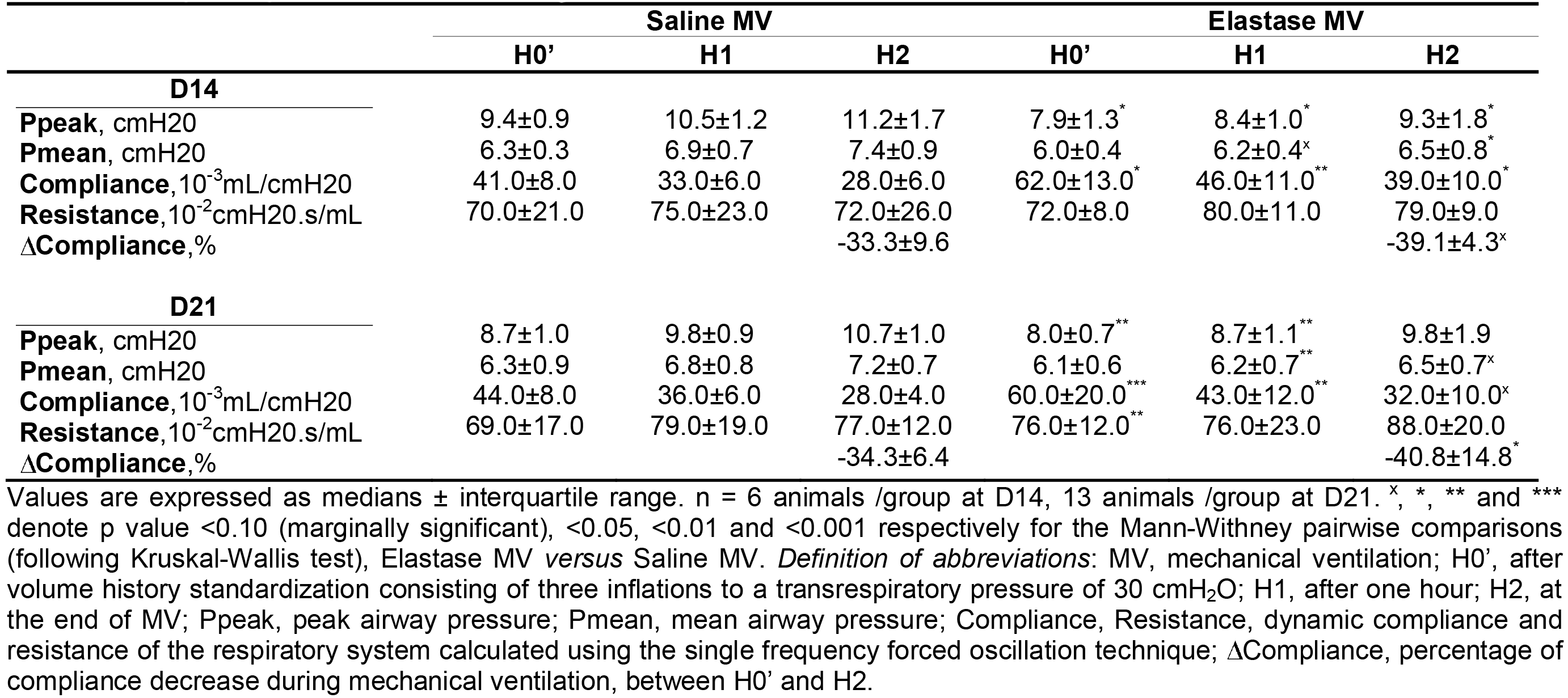
**Respiratory mechanics data** during mechanical ventilation in D14 and D21 Saline and Elastase mice. Values are expressed as medians ± interquartile range. n = 6 animals /group at D14, 13 animals /group at D21. ^x^, *, ** and *** denote p value <0.10 (marginally significant), <0.05, <0.01 and <0.001 respectively for the Mann-Withney pairwise comparisons (following Kruskal-Wallis test), Elastase MV *versus* Saline MV. *Definition of abbreviations*: MV, mechanical ventilation; H0’, after volume history standardization consisting of three inflations to a transrespiratory pressure of 30 cmH_2_O; H1, after one hour; H2, at the end of MV; Ppeak, peak airway pressure; Pmean, mean airway pressure; Compliance, Resistance, dynamic compliance and resistance of the respiratory system calculated using the single frequency forced oscillation technique; ΔCompliance, percentage of compliance decrease during mechanical ventilation, between H0’ and H2.

|  | Saline MV |  |  | Elastase MV |  |  |
| --- | --- | --- | --- | --- | --- | --- |
|  | H0' | H1 | H2 | H0' | H1 | H2 |
| **D14** |  |  |  |  |  |  |
| **Ppeak**, cmH2O | 9.4±0.9 | 10.5±1.2 | 11.2±1.7 | 7.9±1.3<sup>*</sup> | 8.4±1.0<sup>*</sup> | 9.3±1.8<sup>*</sup> |
| **Pmean**, cmH2O | 6.3±0.3 | 6.9±0.7 | 7.4±0.9 | 6.0±0.4 | 6.2±0.4<sup>x</sup> | 6.5±0.8<sup>*</sup> |
| **Compliance**, 10<sup>-3</sup>mL/cmH2O | 41.0±8.0 | 33.0±6.0 | 28.0±6.0 | 62.0±13.0<sup>*</sup> | 46.0±11.0<sup>**</sup> | 39.0±10.0<sup>*</sup> |
| **Resistance**, 10<sup>-2</sup>cmH2O.s/mL | 70.0±21.0 | 75.0±23.0 | 72.0±26.0 | 72.0±8.0 | 80.0±11.0 | 79.0±9.0 |
| **ΔCompliance**, % |  |  | -33.3±9.6 |  |  | -39.1±4.3<sup>x</sup> |
| **D21** |  |  |  |  |  |  |
| **Ppeak**, cmH2O | 8.7±1.0 | 9.8±0.9 | 10.7±1.0 | 8.0±0.7<sup>**</sup> | 8.7±1.1<sup>**</sup> | 9.8±1.9 |
| **Pmean**, cmH2O | 6.3±0.9 | 6.8±0.8 | 7.2±0.7 | 6.1±0.6 | 6.2±0.7<sup>**</sup> | 6.5±0.7<sup>x</sup> |
| **Compliance**, 10<sup>-3</sup>mL/cmH2O | 44.0±8.0 | 36.0±6.0 | 28.0±4.0 | 60.0±20.0<sup>***</sup> | 43.0±12.0<sup>**</sup> | 32.0±10.0<sup>x</sup> |
| **Resistance**, 10<sup>-2</sup>cmH2O.s/mL | 69.0±17.0 | 79.0±19.0 | 77.0±12.0 | 76.0±12.0<sup>**</sup> | 76.0±23.0 | 88.0±20.0 |
| **ΔCompliance**, % |  |  | -34.3±6.4 |  |  | -40.8±14.8<sup>*</sup> |

**Figure 4.**
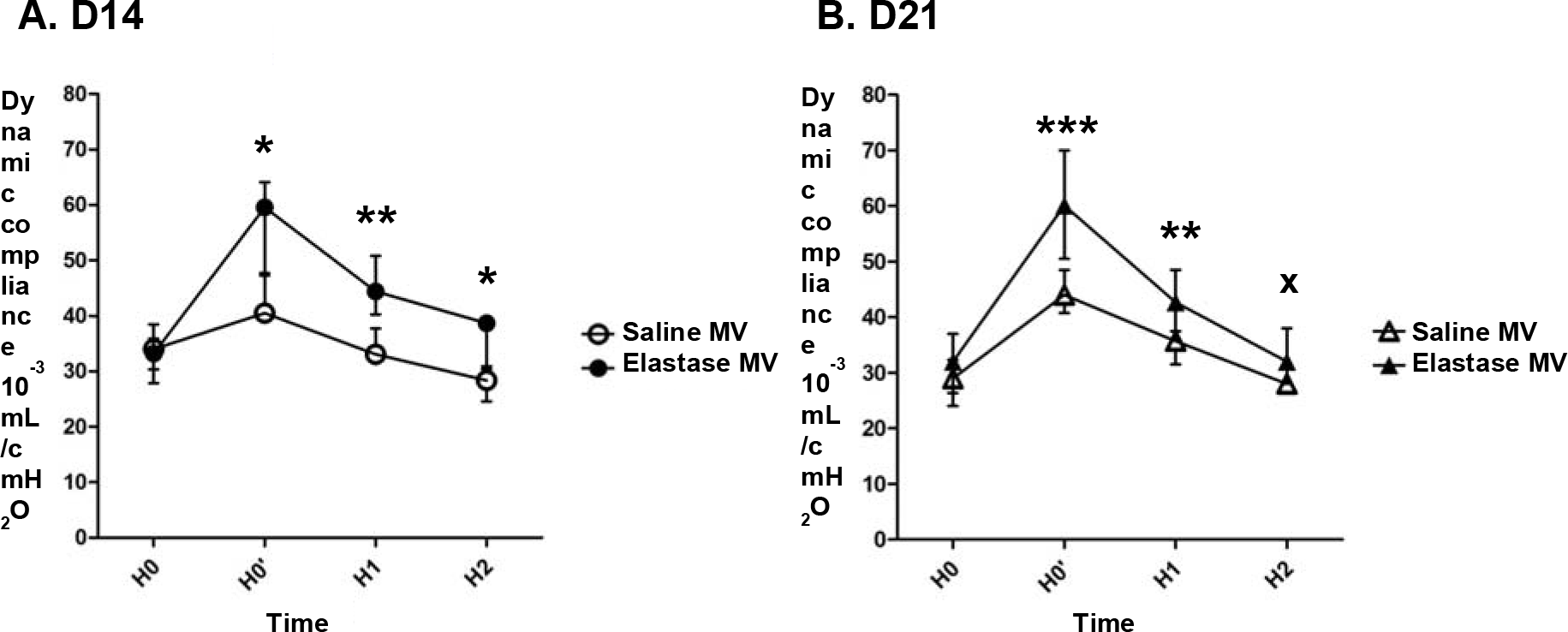
Evolution of dynamic compliance, calculated using the single frequency forced oscillation technique, during two hours of mechanical ventilation in Saline and Elastase mice, at D14 (A) and D21 (B) of tracheal instillation. Values are expressed as median ± interquartile range. n = 6 animals /group at D14, and 13 animals /group at D21. ^±^, *, ** and *** denote p value <0.10 (marginally significant), <0.05, <0.01 and <0.001 respectively for the Mann-Withney pairwise comparisons (following Kruskal-Wallis test), Elastase MV *versus* Saline MV. *Definition of abbreviations*: MV, mechanical ventilation; H0, at ventilator connection; H0’, after volume history standardization; H1, after one hour of MV; H2, at the end of MV.

BAL cellularity was similar between Saline and Elastase mice, at both D14 and D21 (Table S4 in additional file) and differential cell count showed a predominance of macrophages (>90% of total cells) in all groups. Flow cytometric analysis of inflammatory cells from total lung cell suspensions found a marginally significant increase in the percentage of alveolar macrophages in Elastase SV mice compared with Saline SV mice at D14 (Figure 5). These macrophages showed a trend towards an overexpression of CD11b and a stronger autofluorescence, compared with Saline SV mice (Figure 6). Flow cytometric analyses were similar between Elastase SV and Saline SV mice at D21, except for a significant increase in autofluorescence of alveolar macrophages in the former group (Figure S1 and S2 in additional file).

**Figure 5.**
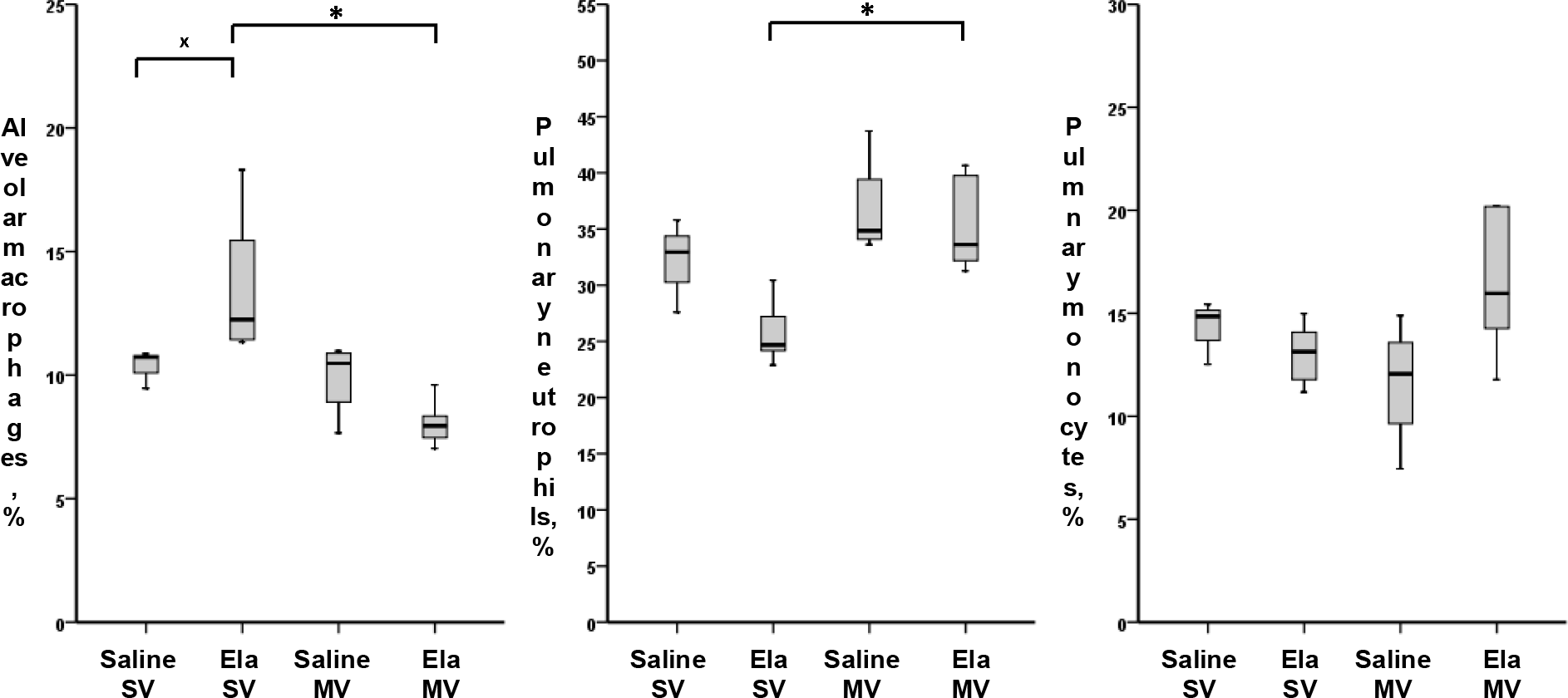
Relative values of pulmonary inflammatory cell populations analyzed by flow cytometry on total lung cell suspensions of mice subjected to spontaneous (SV) or mechanical (MV) ventilation 14 days after instillation of saline (Saline) or elastase (Ela), expressed as percentage of gated cells. Values are expressed as median ± interquartile range. n = 5 animals /group. ^±^ and * denote Benjamini-Hochberg corrected p value <0.10 (marginally significant) and <0.05 respectively for the following Mann-Withney pairwise comparisons (following Kruskal-Wallis test): Saline SV *vs* Saline MV, Saline SV *vs* Ela SV, Ela SV *vs* Ela MV, Saline MV *vs* Ela MV. *Definition of abbreviations*: Ela, Elastase; SV, spontaneous ventilation; MV, mechanical ventilation.

**Figure 6.**
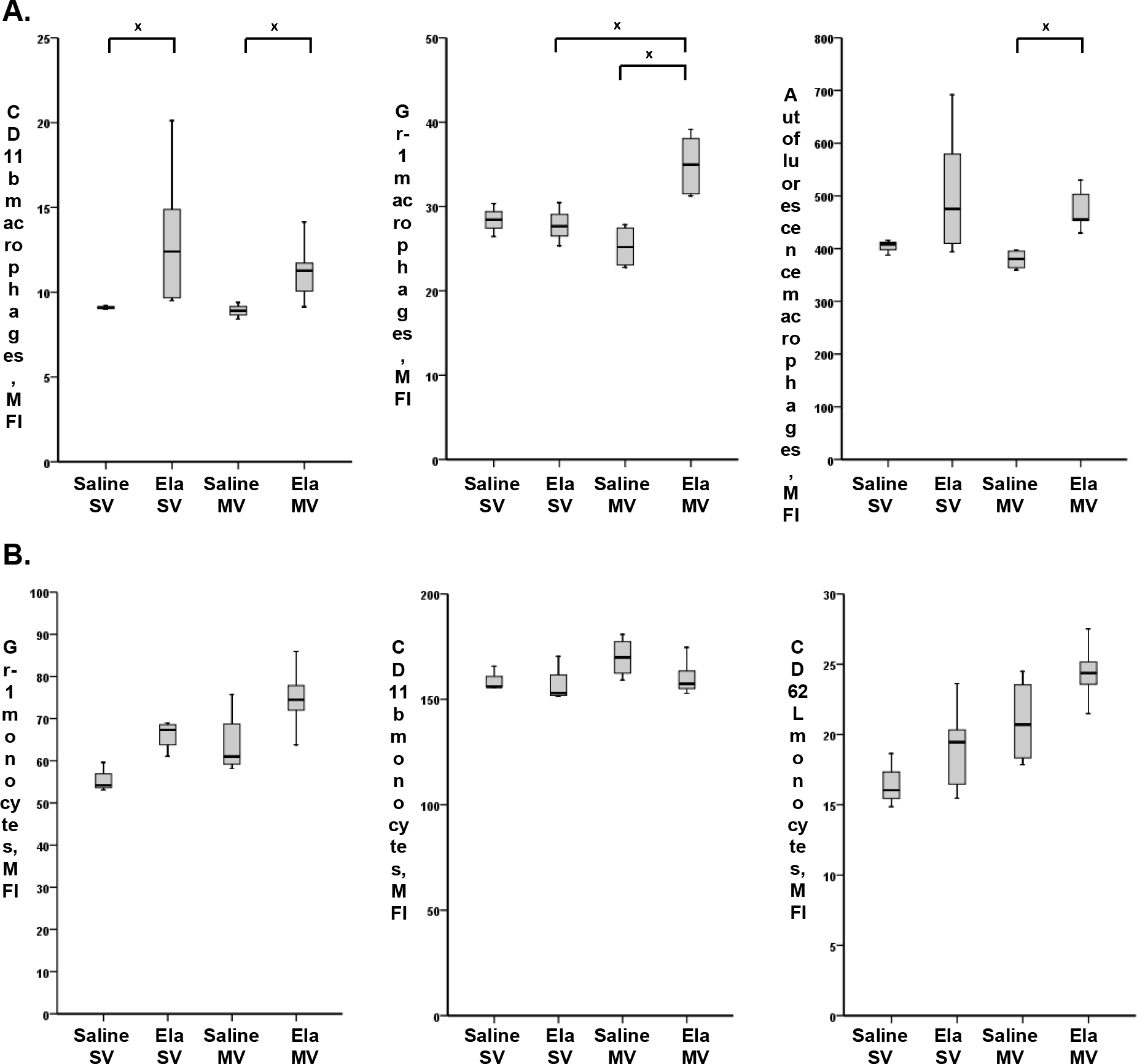
Activation markers of alveolar macrophages (A) and pulmonary monocytes (B), analyzed by flow cytometry on total lung cell suspensions of mice subjected to spontaneous (SV) or mechanical (MV) ventilation 14 days after instillation of saline (Saline) or elastase (Ela), expressed as mean fluorescent intensity (MFI). **A.** CD11b and Gr1 expression, and autofluorescence of alveolar macrophages. **B.** Gr1, CD11b and CD62L expression on pulmonary monocytes. Values are expressed as median ± interquartile range. n = 5 animals /group. ^±^ denotes Benjamini-Hochberg corrected p value <0.10 (marginally significant) for the following Mann-Withney pairwise comparisons (following Kruskal-Wallis test): Saline SV *vs* Saline MV, Saline SV *vs* Ela SV, Ela SV *vs* Ela MV, Saline MV *vs* Ela MV. *Definition of abbreviations*: Ela, Elastase; SV, spontaneous ventilation; MV, mechanical ventilation.

### Effect of mechanical ventilation

A gradual decrease in respiratory system compliance (with associated increase in peak airway pressures) was observed in both Elastase MV and Saline MV mice during the two hours of mechanical ventilation. This decrease was similar between Elastase MV and Saline MV mice at D14, but was greater in the former group at D21 (40.8% *versus* 34.3%, p<0.05, Table 1, Figure 4).

BAL cellularity was comparable between Elastase MV, Elastase SV, Saline MV and Saline SV mice, at both D14 and D21 time points (Table S4 in additional file). Flow cytometric analysis on total lung cell suspensions did not find any significant change in pulmonary inflammatory cell populations or cellular activation state between Saline MV and Saline SV mice, but with a trend towards increased percentage of neutrophils and increased CD62L expression on monocytes in the former group (Figure 5 and 6B).

A significant decrease in the percentage of alveolar macrophages (with a concomitant increase in the percentage of neutrophils) was noted at D14 in Elastase MV mice as compared to Elastase SV mice (Figure 5). This was associated with a change in alveolar macrophages phenotype, with a marginally significant overexpression of Gr1 in the former group (Figure 6A). Pulmonary monocytes also exhibited a modified phenotype, with a maximal expression of CD62L in Elastase MV mice (Figure 6B). A particular stairs aspect was observed in CD62L expression on monocytes, which gradually increased in relation to the successive assaults between Saline SV, Elastase SV, Saline MV and Elastase MV groups. All above mentioned flow cytometry differences were not observed at D21 time point, except for a stronger autofluorescence of alveolar macrophages in Elastase MV mice as compared to Saline MV mice, which was present both at D14 and D21 (Figure S1 and S2 in additional file).

## DISCUSSION

Main results of our study are as follows: i) elastase instillation resulted in a similarly increased mean chord length and respiratory system compliance at D14 and D21, as compared to saline instillation; ii) these alterations were associated with a transient lung infiltration (only at D14) of activated alveolar macrophages CD11b*mid*; iii) lung mechanics was similarly altered during a two hours mechanical ventilation in Elastase and Saline mice, with a gradual decrease in respiratory system compliance; iv) at D14, mechanically ventilated Elastase mice showed a significant increase in the percentage of neutrophils concomitant with a decrease in the percentage of alveolar macrophages in total lung, compared with Elastase animals spontaneously ventilating. Furthermore, alveolar macrophages of mechanically ventilated Elastase mice at D14 overexpressed Gr1, whereas monocytes showed a trend to overexpression of CD62L.

Elastase-induced emphysema model has been described in numerous studies (20–23,27). Murine lungs undergo an intense inflammatory reaction within the first week after elastase instillation, then show a minimal inflammation state in the late phase, after D21. This situation mimics what is observed in human emphysematous lung (17). Moreover, we used a dose of elastase inducing a degree of emphysema similar to that observed after cigarette smoke exposure (near 30% increase in mean chord length) (28). Time course of histological emphysema and alveolar macrophages infiltration observed only at D14 in Elastase SV animals is consistent with previous data from the literature (21–23,27). To our knowledge, our study is the first to provide phenotypic characterization of alveolar macrophages in elastase model, by showing overexpression of CD11b in these cells at D14. These macrophages are similar in their phenotype to pulmonary interstitial macrophages (25,29), but their strong autofluorescence identifies them as alveolar macrophages. CD11b is an adhesion molecule whose expression reflects the level of activation of various inflammatory cells. Activated alveolar macrophages play a central role in the pathophysiology of pulmonary emphysema in mice and humans (30). An increased expression of CD11b has been reported on macrophages collected in induced sputum of COPD patients. Interestingly, CD11b expression intensity was correlated with the severity of airflow limitation (31). High autofluorescence of alveolar macrophages, as observed in Elastase mice (in either SV or MV groups, at both instillation times), has been previously reported in the BAL of smokers (32). A link with tobacco particles phagocytosis has been suggested, without clear biological significance.

We chose normal Vt strategy for its clinical relevance, as Vt close to 8 ml/kg are now widespread in most mechanically ventilated patients (33). In terms of respiratory mechanics, both D14 and D21 Elastase mice responded to mechanical ventilation similarly to Saline mice, decreasing to the same extent their respiratory system compliance within two hours of ventilation. This decrease is reported in various ventilated murine models and results from progressive alveolar derecruitment (34). An identical mechanical response to normal Vt mechanical ventilation has been observed in a TIMP3 KO murine model of emphysema (35). Following two hours of normal Vt mechanical ventilation, we observed a significant increase in the percentage of neutrophils concomitant with decrease in the percentage of alveolar macrophages only in D14 Elastase mice. Neutrophils recruitment to the lung has been already reported as an important mechanism in VILI (11). Cell phenotype changes included Gr1 overexpression by alveolar macrophages, and CD62L overexpression by pulmonary monocytes. Gr1 overexpression is the witness of an activation of macrophages, and has already been demonstrated in infectious circumstances (36). Besides, CD62L is an adhesion molecule whose overexpression on pulmonary monocytes has already been observed during mechanical ventilation and explained by cellular activation related to stretch (12). We postulate that inflammatory response to mechanical ventilation in Elastase mice could be related to pre-existing inflammation reflected by the presence of activated alveolar macrophages CD11b*mid*, and not to altered cellular mechanical properties secondary to parenchymal destruction. Indeed, although morphological and functional lung modifications were similar in Elastase mice at D14 and D21 as compared to Saline animals, no modification in both proportions and activation state of pulmonary inflammatory cells was seen in Elastase mice at D21. Previous data have already demonstrated the early and central role of activation of alveolar macrophages subjected to stretch in the initiation of the inflammatory response to mechanical ventilation (13). Pre-existing macrophage activation could predispose these cells to further activation by mechanical ventilation. Whatever the underlining molecular mechanism, this pulmonary inflammatory response following mechanical ventilation could play a deleterious role in the progression of pulmonary emphysema, through a worsening of lung inflammation level (30,37). Our study had some limitations. First we did not identify any substantial variations in inflammatory cell populations in total lung in Saline mice after normal Vt mechanical ventilation, unlike a previous study with a very close experimental protocol, which highlighted increased number of neutrophils in total lung, and increased expression of CD62L on pulmonary monocytes, in response to normal Vt mechanical ventilation in healthy mice (12). However, we observed a trend towards an increased percentage of neutrophils and expression of CD62L on lung monocytes of Saline MV mice as compared to Saline SV mice. The nature of our control group may explain this lack of significance. Indeed, the Saline SV group (consisting of mice instilled with saline, anesthetized, intubated and maintained in spontaneous ventilation for two hours) was probably subjected to some level of aggression, including pulmonary micro-atelectasis due to hypoventilation. Second, we did not use an infectious challenge (e.g., lipopolysaccharide instillation) in conjunction with elastase instillation and mechanical ventilation. Such a triple hit model may be closer to the frequent clinical scenario of COPD patients requiring mechanical ventilation because of pneumonia, but its implementation and interpretation may be complex.

## CONCLUSION

In conclusion, in a murine model of pulmonary emphysema induced by tracheal instillation of elastase, inflammatory response to normal Vt mechanical ventilation was characterized by an increase in the proportion of pulmonary neutrophils and an activation of alveolar macrophages and pulmonary monocytes. This response was observed only when the emphysema model showed an underlying inflammation, reflected by the presence of activated alveolar macrophages CD11b*mid*.

## LIST OF ABBREVIATIONS

BAL: Bronchoalveolar Lavage

COPD: Chronic Obstructive Lung Disease

FBS: Fetal Bovine Serum

ICU: Intensive Care Unit

MV: Mechanical Ventilation

Pmean: Mean Airway Pressure

Ppeak: Peak Airway Pressure

SV: Spontaneous Ventilation

VILI: Ventilator-Induced Lung Injury

Vt: Tidal Volume

## DECLARATIONS

### Ethics approval

All the experiments were performed in accordance with the official regulations of the French Ministry of Agriculture and the US National Institute of Health guidelines for the experimental use of animals, were approved by the local institutional animal care and use committee, and were conducted in a specific “little animal” plateform (Plateforme Exploration Fonctionnelle du Petit Animal, INSERM U955, Créteil, France).

### Consent for publication

Not applicable

### Availability of data and materials

The datasets during and/or analysed during the current study available from the corresponding author on reasonable request.

### Authors’ contributions

Study concept and design: AR, GV, EG, FR, DI, LB, BM, AMD, JB

Acquisition of data: AR, GV, EG, FR, JTVN

Analysis and interpretation of data: AR, GV, EG, FR, JTVN, BM, AMD, JB

Drafting of the manuscript: AR, GV, EG, AMD, JB

Critical revision of the manuscript for important intellectual content: all authors

Statistical analysis: AR, GV, AMD

Study supervision: GV, JB

All authors read and approved the final manuscript.

## Acknowledgments

We are very grateful to the following members of U955 INSERM unit for their scientific and technical assistance: Stéphane Kerbrat, Dr Sabine Le Gouvello, Dr Laurent Boyer (Equipe 4, INSERM U955, Créteil), Rachid Souktani (Plateforme Exploration Fonctionnelle du Petit Animal, INSERM U955, Créteil), Aurélie Guguin and Adeline Henri (Plateforme Cytométrie en Flux, INSERM U955, Créteil), Xavier Decrouy (Plateforme Imagerie, INSERM U955 Créteil).

